# GABA antagonises ABA perception and signalling in plants

**DOI:** 10.64898/2026.05.14.725281

**Authors:** Cui-Zhu Feng, Yun-Xin Luo, Hui Zhou, Ping Lin, Bin-Rui Liu, Liangyu Guo, Aolin Jia, Xin Zhang, Yun Zhou, Chun-Peng Song, Yu Long

## Abstract

Abscisic acid (ABA) is a central regulator of plant growth and stress responses. However, it also negatively impact germination, growth, and reproductive performance, highlighting the need for mechanisms that fine-tune its signalling. Here, we identify γ-aminobutyric acid (GABA) as an endogenous antagonist of plant ABA receptor. We show that GABA binds the clade A protein phosphatase–ABI1, disrupting its association with the ABA receptor PYL1, blocking ABA binding, thereby attenuating ABA signalling. GABA modulates ABA-control of stomatal aperture and seed germination, demonstrating its broad physiological influence. Notably, GABA accumulation during stress coincides with the subsequent decline in ABA, consistent with a delayed feedback signal of ABA activity. Together, this work identifies GABA as an antagonist of ABA and provides a mechanism balancing plant stress resilience and growth.

## Introduction

Global climate change, particularly the increasing frequency and intensity of abiotic stress, poses major challenges to agriculture. Crop plants must not only withstand high levels of stress but also maintain growth and productivity under these adverse conditions^1,2^. Abscisic acid (ABA) is a central plant hormone that orchestrates abiotic stress responses while also regulating growth^3,4^. The core ABA signalling module comprises PYRABACTIN RESISTANCE/PYR1-LIKE/REGULATORY COMPONENT OF ABA RECEPTOR (PYR/PYL/RCAR) receptors; clade A type 2C protein phosphatases (PP2Cs), such as ABSCISIC ACID INSENSITIVE 1 (ABI1); and SUCROSE NON FERMENTING (SNF1)-related kinases (SnRKs), such as SnRK2.6 (Open Stomata 1, OST1)^5–7^. In this pathway, ABA promotes the interaction of PYR/PYL/RCARs with PP2Cs, inhibiting PP2C phosphatase activity. This inhibition releases SnRKs from repression, enabling them to phosphorylate downstream effectors and trigger stress-adaptive responses^8–11^.

However, ABA signalling also carries costs. Prolonged, excessive ABA response can impose a secondary stress on plants, through elevated reactive oxygen species (ROS) accumulation and sustained stomatal closure^12^. Thus, terminating ABA signalling is as important as activating it, to fine-tune the balance between stress resistance and growth. Plants suppress ABA signalling through two main mechanisms: restricting ABA accumulation and repressing downstream signalling components^13,14^. Synthetic ABA antagonists have been developed to inhibit ABA signalling and regulate plant growth^15–18^; however, no endogenous antagonist of ABA receptors has been identified.

γ-Aminobutyric acid (GABA, a non-protein amino acid, is an ancient signalling molecule conserved across biological kingdoms^19^. In animals, GABA functions as a major inhibitory neurotransmitter, dampening excitatory signalling and promoting relaxation under stress^20^. In plants, GABA accumulates under diverse abiotic and biotic stresses and has been described as a stress-related signalling metabolite^21,22^. GABA is synthesised by glutamate decarboxylase (GAD), with some isoforms activated by Ca^2+^/calmodulin binding to a C-terminal autoinhibitory domain and by cytosolic acidification, two of the earliest cellular responses to stress^23^ Accordingly, GABA accumulation is widely recognised as a metabolic marker of plant stress^21,24^.

Over the past decade, exogenous GABA has been reported to enhance stress resistance in multiple plant species, though the underlying mechanisms are far from fully described^24^. We previously showed that GABA reduces stomatal opening in response to light by inhibiting ALUMINUM-ACTIVATED MALATE TRANSPORTER 9 (ALMT9)^19^, but how GABA antagonises ABA-driven stomatal closure and broader ABA signalling has remained unresolved until now. Here, we demonstrate that GABA directly antagonises ABA signalling by blocking the binding between ABA and PYL1–ABI1 co-receptors. This antagonism goes well beyond regulating stomatal aperture by affecting multiple processes, including gene expression, seed germination, root growth, and stomatal regulation. Our findings identify GABA as the first endogenous ABA receptor antagonist, providing a mechanistic basis for how GABA contributes to plant stress resilience through modulation of ABA perception and signalling.

## Results

### GABA blocks the ABA signal *in vitro*

To explore whether GABA interacts with ABA signalling, we expressed *PYL1*, *ABI1*, *OST1*, and *SLOW ANION CHANNEL ASSOCIATED 1* (*SLAC1*) in *Xenopus laevis* oocytes to reconstitute the canonical ABA signal pathway that initiates stomatal closure in *Arabidopsis thaliana*^25^. As expected, stepwise increases in ABA application increased SLAC1 S-type anion channel activity, as detected by a two-electrode-voltage-clamp (TEVC) assay, confirmed that this system was suitable for assessing ABA signalling (**Figure 1A**). To measure the impact of GABA on ABA signal transduction, we injected GABA into the *PYL1/ABI1/OST1/SLAC1-*expressing oocytes in increasing amounts. Both GABA and muscimol, the potent GABA analogue and mammalian GABA_A_ receptor agonist^26^, reduced the SLAC1 S-type anion channel activity induced by 10 μM ABA (**Figure 1B and 1C**). Inhibition of SLAC1 activity increased with increasing GABA concentration (**Figure 1B**), whereas muscimol inhibited SLAC1 near completely at all concentrations tested (**Figure. 1C**).

**Figure 1.**
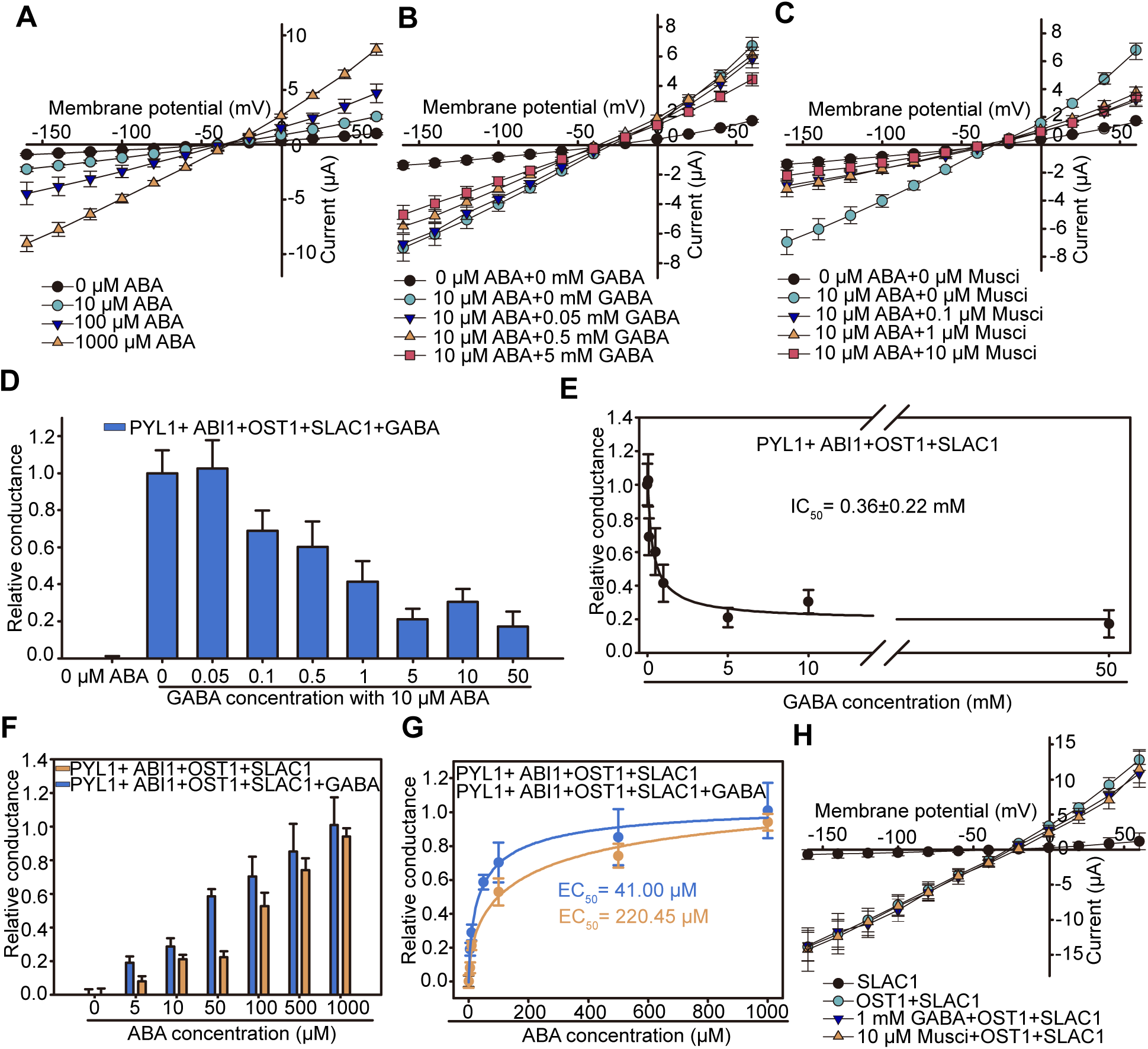
GABA blocks canonical ABA signal transduction via the PYL1/ABI1/OST1/SLAC1 pathway. **A.** ABA effect on SLAC1 S-type anion channel activity in *X. laevis* oocytes injected with *PYL1*, *ABI1*, *OST1*, and *SLAC1* cRNA. **B.** GABA effect on ABA-induced SLAC1 channel activity in *X. laevis* oocytes expressing system in (**A**). **C.** Effect of the GABA analogue muscimol (Musci) on ABA-induced SLAC1 channel activity in *X. laevis* oocytes expressing system in (**A)**. **D.** Effect of increasing GABA concentration on ABA-induced SLAC1 activation. **E.** Kinetic analysis of ABA-activated SLAC1 conductance at increasing GABA concentrations. Data are fitted with the Hill equation. **F, G.** Effect of increasing ABA concentrations on GABA blockage of ABA-activated SLAC1 activity (GABA at 1mM). Data are fitted with the Hill equation. **H.** Effect of GABA and muscimol on OST1 and SLAC1 activity in *X. laevis* oocytes. For **A-H**, data are mean ± s.e. (*n* = 12). Values of ABA and GABA are the calculated final concentrations in oocytes of the injected substrate.

More detailed kinetic analysis of PYL1/ABI1/OST1-mediated SLAC1 activation by 10 μM ABA indicated the half-maximal inhibition (IC_50_) by GABA was 0.36 ± 0.22 mM (**Figure. 1D and 1E**). Increasing ABA concentrations in the presence of 1 mM GABA revealed that the half-activating ABA concentration for SLAC1 (EC_50_) was increased five times by GABA (**Figure. 1F and 1G**). Removing PYL1 and ABI1 from the assay, which allowed OST1 to directly activate SLAC1, abolished GABA inhibition of SLAC1 activity completely (**Figure. 1H**), indicated that GABA blockage occurred upstream of OST1 and SLAC1, most likely via ABI1 and/or PYL1.

### GABA blocks ABA-induced stomatal closure

To test whether GABA inhibition of ABA-induced SLAC1 activation occurred *in vivo*, we used patch clamp electrophysiology on guard cell protoplasts isolated from *Arabidopsis* leaves. Consistent with the oocyte assays, GABA suppressed ABA-induced S-type anion currents in protoplasts (**Figure. 2A and 2B**). Our previous work showed that exogenous GABA alone does not stimulate a change in stomatal aperture but rather it modulates the extent of opening^19^; here, we show it can partially inhibit ABA-induced stomatal closure (**Figure. 2C**). Collectively, our findings so far suggest an antagonistic role of GABA in ABA signalling *in planta* that we further explored using endogenous manipulation of GABA.

**Figure 2.**
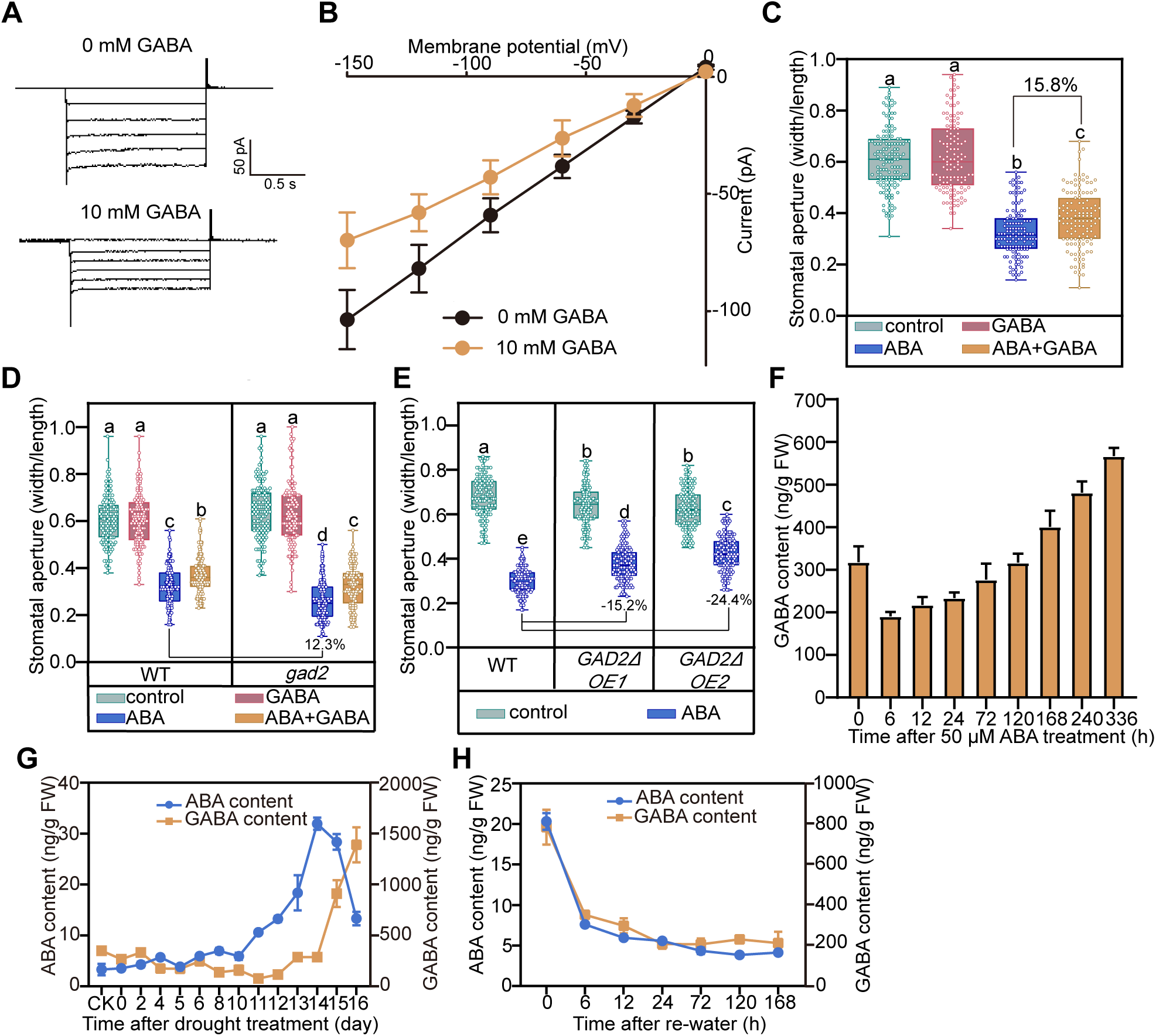
GABA regulates ABA-induced S-type anion current activation and stomatal closure, and accumulates when ABA declines during drought. **A,B.** Whole cell recording (**A**) and steady-state current–voltage relationships (**B**) for ABA-induced slow type anion current in guard cell protoplasts with or without GABA treatments. Data are mean ± s.e. (*n = 6*). **C.** Wild-type stomatal aperture in response to treatments in epidermal peels. **D,E.** Stomatal aperture of wild-type compared with the *gad2* mutant (**D**) and *GAD2*Δ *OE* lines (**E**) in response to treatments on epidermal peels. **F.** GABA accumulation following ABA induction in wild-type seedings. **G,H.** ABA and GABA contents during drought stress (**G**) and recovery (**H**). For **c**–**e**, box and whisker plots show interquartile range and extremes (*n* > 120 for each material and treatment). Letters indicate significant differences at *P* < 0.05 by ANOVA with Tukey’s comparison test. Percentages indicate GABA (**C**) or *GAD2* gene expressing alteration (**D** and **E**) caused influence to stomatal closure compared with ABA treated or wild-type control. For **F**–**H**, data are mean ± s.e. (*n* = 3). FW, fresh weight; CK, none drought treated control.

The *gad2* mutant has very low leaf GABA content^19^. ABA was more effective in closing *gad2* stomata than for wild-type, but ABA-induced stomatal closure in both genotypes was significantly attenuated by addition of GABA (**Figure 2D**). Supporting this finding, increasing endogenous GABA content in leaves, by ubiquitous expression of the constitutively active *GAD2*Δ (*GAD2*Δ OE; **Figure S1A and S1B**), resulted in guard cells being less responsive to ABA (**Figure 2E**). However, as increased GABA content has been shown to enhance stress resilience of plants, including drought tolerance (**Figure S1C and S1D**)^19,23^, our findings raise a key question: how can GABA promote stress tolerance if it antagonises ABA-induced stomatal closure?

To investigate this, we analysed the relationship between ABA and GABA accumulation. We first observed that ABA itself gradually induced an increase in GABA concentration (**Figure 2F**). Following the onset of drought, ABA concentration rose dramatically, followed by GABA accumulation a few days later when ABA content started to decline (**Figure 2G**). After re-watering, both ABA and GABA concentration decreased similarly (**Figure 2H**). A similar pattern was observed under osmotic and salt stress, where ABA accumulated first, and GABA accumulated at later stages synchronised with declining ABA (**Figure S2A–D**). In contrast, ABA accumulation did not occur following GABA treatment (**Figure S2E**), nor was it affected by endogenous GABA concentrations following osmotic stress (**Figure S2F**). We propose that GABA functions as an antagonist of ABA signalling, but not immediately after ABA accumulation or when ABA levels are high — conditions under which GABA is less effective (e.g., **Figure 1G**). Instead, GABA acts during later stages of stress, attenuating the ABA signal when ABA concentration is in decline.

### GABA broadly suppresses ABA-mediated processes

To test whether GABA antagonism extends beyond stomatal regulation, we examined its effects on other ABA-related traits. First, we tested if GABA inhibited transcription of ABA-induced gene expression, using the classic ABA transcriptional activated gene *RESPONSIVE TO DESICCATION 29A* (*RD29A*) promoter as a reporter. We built an ABA signal pathway in mesophyll protoplasts by transiently expressing *PYL1*, *ABI1*, *OST1*, *ABSCISIC ACID–RESPONSIVE ELEMENT–BINDING FACTOR 2* (*ABF2*; activated by OST1 to induce *RD29A* expression), luciferase under control of the *RD29A* promoter (*ProRD29A:LUC*), and *35S:Ren* (Renilla) served as an internal control. Dual-luciferase assays showed that ABA induced the *ProRD29A:LUC* signal, which was attenuated by GABA; GABA by itself did not cause any change in LUC signal (**Figure 3A**). We verified that GABA antagonised ABA-induced *RD29A* and *RD29B* genes expression, as ABA-response marker genes^2^, using quantitative PCR in *Arabidopsis* seedings (**Figure 3B**). Further, ABA-induced gene expression increased in mutant lines with low endogenous GABA (*gad1*, *gad2*, and *gad1245*), and decreased in the GABA overexpression lines (**Figure 3C and 3D**). These results indicate that GABA antagonises broad ABA signalling, so we explored further physiological contexts where GABA may affect ABA-induced plant responses.

**Figure 3.**
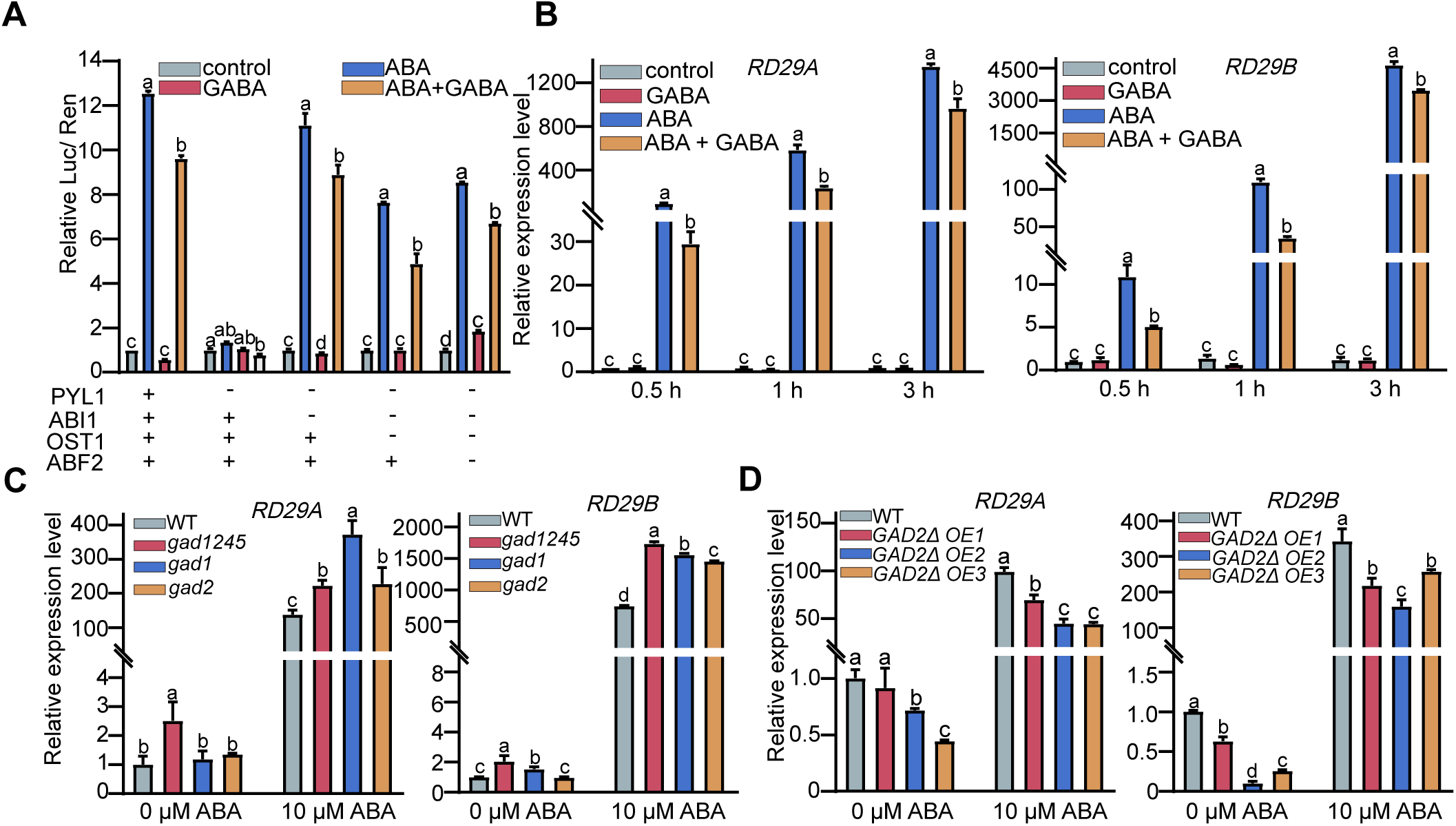
GABA blocks ABA-induced gene expression. **A**. Effect of GABA on ABA-induced *RD29A* promoter activity after transient transfection by different gene expression constructs in *Arabidopsis* mesophyll protoplasts. **B.** RT-qPCR analysis of *RD29A* and *RD29B* expression at different time points after hormone treatment. **C,D.** RT-qPCR analysis of *RD29A* and *RD29B* expression in *gad* mutants (**C**) and *GAD2Δ OE* (**D**) lines after 1 h of ABA treatment. For **B**–**D**, data are mean ± s.e. (*n* = 3). Letters indicate significant differences at *P* < 0.05 by one-way ANOVA test.

Seed dormancy and inhibition of root elongation are two key traits influenced by ABA. GABA itself had no impact on seed germination, but concentrations ≥0.5 mM GABA decreased the dormancy imposed by both 0.3 and 0.5 μM ABA (**Figure 4A and 4B**). Muscimol again proved a powerful antagonist of ABA, with the impact of 10 μM muscimol on ABA-induced dormancy being similar to 2–5 mM GABA (**Figure 4A and 4B**). Consistent with these findings, alteration of endogenous GABA through *gad* knockout mutants resulted in lower germination rates in the presence of ABA, while *GAD2*Δ OE increased seed germination (**Figure 4C–F**). GABA also reduced root growth inhibition by ABA. Again, GABA had no effect by itself on root growth but at concentrations ≥0.5 mM GABA, could antagonise root growth reductions caused by up to 10 μM ABA. 10 μM muscimol was effective in alleviating inhibition at both 5 and 10 μM ABA (**Figure S3**). Collectively, our results demonstrate that both exogenous and endogenous GABA is a powerful antagonist to a range of ABA-induced genetic and physiological processes in plants.

**Figure 4.**
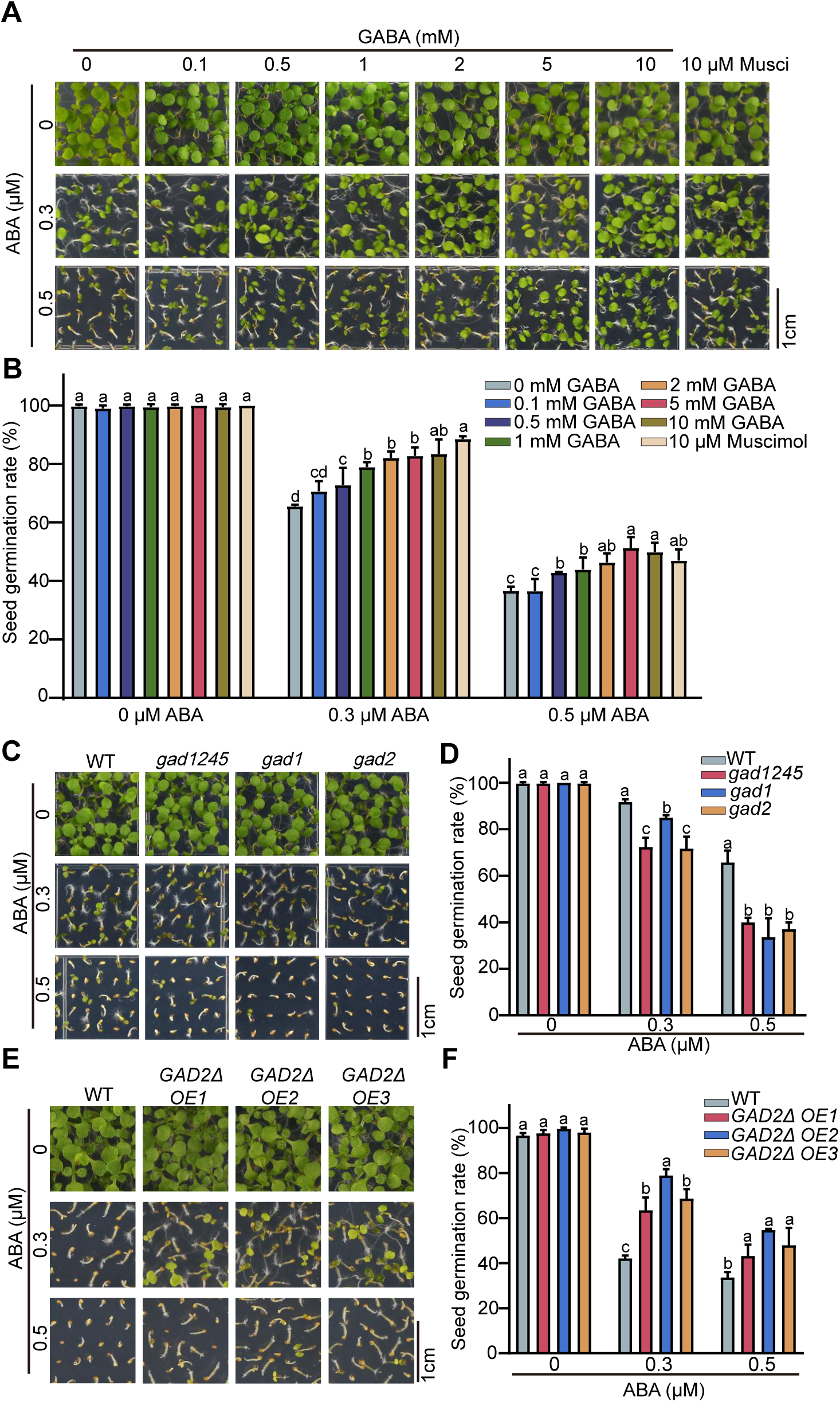
GABA blocks ABA-inhibition of seed germination. **A,B**. Seed germination of wild-type (WT) *Arabidopsis* under different treatments. **C,D.** Seed germination of WT and *gad* mutants under different ABA treatments. **E**,**F.** Seed germination of WT and *GAD2Δ OE* lines under different ABA treatments. For **B**, **D**, and **F**, data are mean ± s.e. (*n* ≥ 100). Letters indicate significant differences at *P* < 0.05 by one-way ANOVA test.

### GABA blocks ABA–PYL1–ABI1 formation

Our results have indicated that GABA is likely to act directly on either ABI1, PYL1, or both (**Figure 1D–H**), so we investigated this mechanism further. The initial event in ABA signal transduction is ABA binding to the PYR/PYL/RCAR receptor (PYL1), which recruits PP2C (ABI1) to the ABA–PYR/PYL/RCAR complex. Therefore we tested whether GABA blocks the ABA-induced PYL1–ABI1 interaction. A pull-down assay confirmed that ABA induced the PYL1–ABI1 interaction, which was blocked by increasing concentrations of GABA, or muscimol (**Figure 5A**). A yeast 2-hybrid assay confirmed that GABA weakened the ABA-induced PYL1–ABI1 interaction (**Figure S4A**), further confirmed through split-luciferase and bimolecular fluorescence complementation assays in *Nicotiana benthamiana* leaves that showed that increased concentration of endogenous GABA reduced the PYL1–ABI1 interaction (**Figure 5B; Figure S4B**). All of these data are consistent with GABA inhibiting the ABA-induced PYL1–ABI1 interaction to block ABA signal transduction.

**Figure 5.**
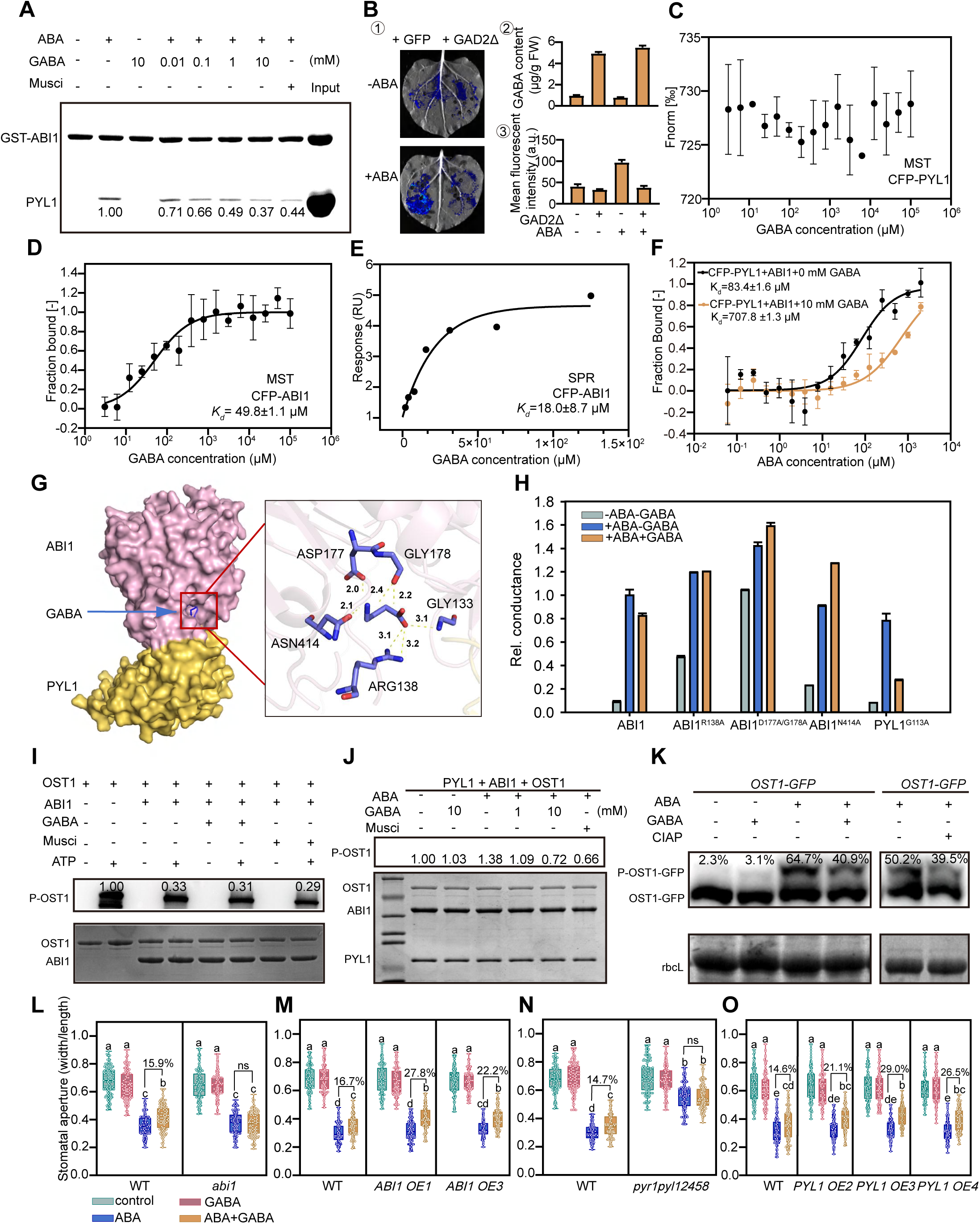
GABA binds ABI1 and blocks the ABA-induced PYL1–ABI1 co-receptor association. **A.** *In vitro* GST pull down assay, with immobilised ABI1, under different treatments. Numbers give protein band intensity normalised to ABA without GABA treatment (lane 2). **B.** *In vivo* split-luciferase assays on ABI1-LUC^N^ and PYL1-LUC^C^ interaction. GABA content and fluorescence intensity are in ② and ③. Data are mean ± s.e. (*n* = 3). **C**,**D.** Binding between GABA and CFP-PYL1 (**C**) or CFP-ABI1 (**D**) analysed using MicroScale Thermophoresis (MST). Data are mean ± s.e. (*n* = 3). **E.** Binding between GABA and CFP-ABI1 analysed using Surface Plasmon Resonance (SPR). **F.** MST assay for binding between ABA and PYL1-ABI1 ± GABA. Data are mean ± s.e. (*n* = 3). **G.** Molecular docking for the binding between GABA and ABI1. **H**. Effect of GABA on ABA-induced SLAC1 activation with different ABI1 mutants. Data are mean ± s.e. (*n* = 12). **I,J.** *In vitro* phosphatase assay showing GABA impact on ABI1 activity in absence (**I**) and presence (**J**) of PYL1 and ABA. OST1 autophosphorylation (P-OST1) was used as the phosphatase activity reporter. Numbers in (**i**) give band intensity normalised to the no ABA/no GABA control (lane 1). **K.** *In vivo* phosphatase assay for OST1-GFP phosphorylation under different treatments. Numbers give phosphorylated (P-OST1-GFP):unphosphorylated (OST1-GFP) ratios. **L–O**, GABA effect on ABA-induced stomatal closure in *abi1* (**L**), *ABI1 OE* (**M**), *pyr1pyl12458* (**N**), and *PYL1 OE* (**O**) lines. Box and whisker plots show interquartile range and extremes (*n* > 120). Letters indicate significant differences at *P* < 0.05 by ANOVA with Tukey’s comparison test. FW, fresh weight; a.u., arbitrary units; ns, not significant; Fnorm, the normalized fluorescence; CIAP, calf intestine alkaline phosphatase.

To further resolve this mechanism, we tested whether GABA binds to PYL1 and/or ABI1 using microscale thermophoresis (MST) assays. PYL1 showed no combination with GABA (**Figure 5C**), whereas binding between ABI1 and GABA was observed with a dissociation constant (*K_d_*) of 49.8 ± 1.1 μM (**Figure 5D**), and 1.0 ± 1.2 μM for muscimol (**Figure S5A**). Surface Plasmon Resonance (SPR) confirmed that binding could occur in the same order of magnitude (*K_d_*= 18.0 ± 8.7 μM; **Figure 5E**), with zero interaction detected GABA and Cyan Fluorescent Protein (CFP), the non-binding control (**Figure S5C and S5D**). Further MST assays demonstrated that GABA alone did not influence the binding affinity between PYL1 and ABA (**Figure S6**). In the presence of ABI1 however, GABA significantly decreased the binding affinity of the PYL1 for ABA, increasing the *K_d_* from 83.4 µM to 707.8 µM (**Figure 5F**).

Next, we utilized molecular docking to predict the structural model of PYL1–ABI1 in complex with GABA. The model with most reliability indicated that GABA binds at the interaction interface between PYL1 and ABI1 with 3 and 1 amino acid binding in ABI1 and PYL1 respectively (**Figure 5G**). To validate the prediction, we mutated these residues to alanine (ABI1^R138A^, ABI1^D177A/G178A^, ABI1^N414A^, and PYL1^G113A^) and examined their effects using the heterologous ABA pathway assay in *Xenopus* oocytes (**Figure 1**). With TEVC recoding on SLAC1-mediated anion conductance, we found that the mutations at 3 sites in ABI1 abolished the blocking of GABA to ABA signaling, These results indicate that GABA binds with ABI1 and blocks PYL1-ABI1 interaction at their interaction interface and thus suppresses antagonizes ABA-PYL1-ABI1 interaction. Collectively, these results support a model in which GABA antagonise ABA–PYL1–ABI1 complex formation.

We then explored how this antagonism might impact ABA signalling. ABI1 regulates ABA signalling by controlling SnRK kinase activity; for stomatal closure, this materialises, in part, through upregulation of ABI1 inhibition of OST1 autophosphorylation^5–8^. To test whether GABA directly affects ABI1 phosphatase activity, we performed *in vitro* phosphatase assays that showed that GABA had no direct effect on ABI1 activity, with OST1 autophosphorylation remaining unchanged in the presence or absence of GABA (**Figure 5I**). However, by incorporating PYL1 and ABA into the assay system, ABA signalling could enhance OST1 autophosphorylation, whereas GABA progressively suppressed this enhancement in a concentration-dependent manner (**Figure 5J**). We conclude that both PYL1 and ABI1 are required for GABA antagonism; specifically, GABA disrupts the PYL1–ABI1 interaction, blocking ABA signalling and ABA-dependent inhibition of ABI1 (**Figure 5J**). Consistent with these *in vitro* findings, *in vivo* OST1 phosphatase assays confirmed that ABA induced OST1 phosphorylation, whereas GABA attenuated this response, with CIAP (Calf Intestine AlkalinePhosphatase) reduced the OST1-phosphorylation as negative control (**Figure. 5K**).

We used genetic analyses to test and validate our model. ABA-induced stomatal closure was analysed in the *abi1-3* mutant^27^, *ABI1* overexpression (OE) lines, the *pyr1pyl12458* sextuple mutant^28^, and *PYL1* OE lines in the presence of GABA. The *abi1-3* mutant was insensitive to GABA (**Figure 5L**), whereas *ABI1* OE lines showed stronger inhibition of ABA-induced stomatal closure by GABA (**Figure 5M**). Similarly, the *pyr1pyl12458* sextuple mutant was insensitive to GABA (**Figure 5N**), while PYL1 OE lines were more sensitive (**Figure 5O**). For all genotypes, darkness was used as the control stimulus for stomatal closure. GABA by itself could inhibit dark-induced closure in all of these lines, indicating that GABA exerts distinct effects on closure responses mediated by darkness independent of ABA signalling (**Figure S7**). Seed germination assays further supported this conclusion, where normal ABA inhibition of germination was not relieved by GABA in *abi1-3* or *pyr1pyl12458* mutants. By contrast, GABA more effectively counteracted ABA-induced dormancy in *ABI1* OE and *PYL1* OE lines (**Figure S8**).

Taken together, these results support a model in which GABA binds ABI1, disrupts ABI1–PYL1 interaction, prevents ABA from stabilising the PYL1–ABI1 complex, and thereby reduces ABI1 phosphatase activity to antagonise ABA signalling. As GABA directly targets ABI1, a core component of the ABA co-receptor complex, we propose that GABA functions as an ABA receptor antagonist.

## Discussion

Our findings identify a previously unrecognised role for GABA as an endogenous ABA receptor antagonist in plants, summarised in our simplified model (**Figure 6**). Mechanistically, GABA binds ABI1 and disrupts its interaction with PYL1, thereby preventing co-receptor assembly. This inhibition ultimately attenuates ABA signalling, establishing GABA as a physiological inhibitor of ABA receptor function.

**Figure 6.**
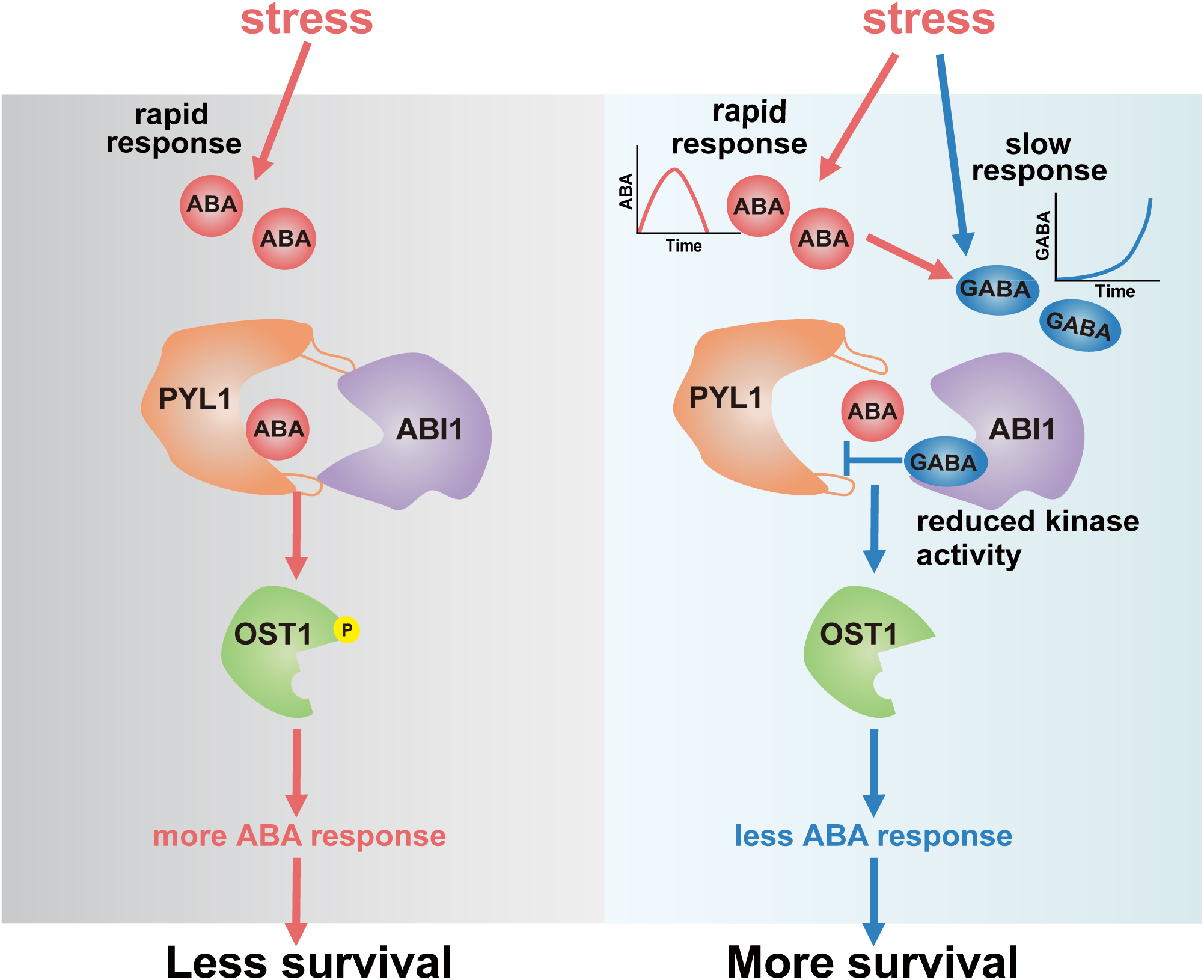
Working model for GABA as an ABA antagonist. When plants face stress, ABA rapidly accumulates; in turn, ABA induces GABA accumulation as a secondary, and delayed modulator. GABA binds to ABI1, which impairs formation of the ABA-induced ABI1–PYL1 signalling complex, thereby suppressing ABA signal transduction and ultimately enhancing plant stress tolerance.

We propose that GABA functions not as a rapid stress-responsive signal, when ABA concentrations are at their highest, but as a slower, modulatory signal that contributes to stress recovery. This conclusion stems from the findings that ABA and GABA concentration increases are asynchronous and of a concentration-dependency in the antagonism between GABA and ABA. GABA binds ABI1 with high affinity (*K_d_* = 18 µM in SPR and 50 µM in MST assays), but physically inhibits ABA signalling only at higher concentrations (∼1 mM). In our experiments, we used 1–10 µM ABA — above typical intracellular levels — to elicit clear ABA-dependent phenotypes and biochemical responses, while applying 1 mM GABA, which approximates endogenous concentrations in plant cells^29^. By contrast, muscimol, the potent GABA analogue, blocked ABA responses at much lower concentrations, further supporting the concentration dependency of both GABA and ABA in manifesting an antagonism. The finding that *almt12* knockout disrupted GABA inhibition of dark-induced stomatal closure (**Figure S7E**) without eliminating GABA inhibition of ABA signalling supports the notion that GABA acts on multiple targets, including ALMT channels¹L and the ABI1–PYL1 complex, and that ALMTs are not required for GABA-mediated suppression of ABA signalling.

Physiologically, maintaining low ABA activity is critical for normal growth. We propose that GABA may contribute to this regulation, which is consistent with the considerable literature indicating GABA improves plant survival and resilience following stress^21,22,24^. We propose that the concentration-dependent antagonism of GABA, together with its gradual accumulation, provides an important buffering mechanism under stress. While ABA acts as a rapid, strong stress signal, GABA moderates ABA impact, preventing excessive or prolonged ABA activity and thereby balancing growth with stress resistance.

GABA is best known as an inhibitory neurotransmitter and modulator of relaxation in animals^20^. Here, we propose that, in plants, GABA also functions as a modulatory signal during stress but via a pathway not found in animals, through attenuation of ABA signalling to prevent detrimental long-term impacts of stress. Unlike previously described ABA antagonists, which are synthetic, costly, and environmentally persistent ^15–18^, GABA is the first identified endogenous ABA inhibitor. Its natural abundance, low cost, and non-hazardous properties make it promising both for field applications as a chemical treatment, and for manipulation through molecular breeding. In addition, while gibberellic acid (GA) has been described as a natural ABA antagonist^30^, its mode of action is different and indirect, blocking downstream signal transduction^31^. While GA and ABA mutually inhibit the accumulation of each other^31^, we show that GABA accumulation is induced by ABA; whereas GA content reduces in response to stress, GABA accumulates, indicating its greater potential for assisting in plant stress adaptation.

In summary, our findings establish that GABA is an endogenous antagonist of an ABA receptor, blocking ABA perception and signalling transduction in plants. The delayed accumulation of GABA relative to ABA during stress supports its role as a secondary signal that tempers ABA activity, preventing excessive responses and assisting in the balance of growth regulation and stress adaptation. Beyond providing a mechanistic explanation for the long-standing observations of GABA’s effects on plant physiology, this work reveals a metabolite-based layer of hormone regulation, and identifies a natural ABA antagonist with potential for crop improvement.

## Method details

### Plant materials and growth

All experiments were performed on *Arabidopsis* (*Arabidopsis thaliana*) Columbia-0 (Col-0) lines. The *gad1*, *gad2*, *gad1245*, *abi1-3*, and *pyr1pyl12458* knockout mutants, *proSuper:OST1-GFP* plants and constitutively active *GAD2*Δ mutant, were described previously^19,27,32–34^ For stable transformations of *GAD2*Δ, *PYL1*, and *ABI1* overexpression (OE) lines, coding sequences of these genes were amplified separately by PCR with the primer listed in Supplementary Table 1 and generated by Gibson assembly into pCAMBIA1300 driven with *35S* promoter, and then introduced into wild-type plants using Agrobacterium (*Agrobacterium tumefaciens*) strain GV3101 respectively^35^. Plants were grown in long-day conditions (120 μmol·m^-2^·s^-1^, 16 h light/8 h darkness, 22°C/18°C).

### Electrophysiology assays

Two-electrode voltage-clamp (TEVC) assays in *Xenopus laevis* oocytes were used to measure SLAC1 anion channel activity. Full-length coding sequences of *PYL1*, *ABI1*, *OST1-YFP^N^* (N-terminal domain of Yellow Fluorescent Protein) and *SLAC1-YFP^C^* (C-terminal domain of YFP) were cloned into the oocyte expression vector pGXBG by Gibson assembly. All primers are listed in Supplementary Table 1. Harvest of complementary RNA (cRNA), oocyte preparation for cRNA injection, and whole oocyte TEVC were performed as previously described^36^.

For ABA and GABA/muscimol treatments, 50 nL of treatment or a water control were pre-injected into oocytes 1 h before the TEVC recording. Based on an average oocyte volume of 500 nL, the introduced concentration in oocytes is expected to be 10 times less than the injected concentration^37^; figure legends give physiological concentrations. Whole-oocytes ionic currents were recorded with an OC-725C amplifier (Warner) and digitised using an Axon Instruments Digidata 1550B low-noise data acquisition system (Molecular Devices) controlled by pClamp acquisition software (Molecular Devices). TEVC recordings were performed with 2000 × gain, 20 kHz acquisition rate, and 500 Hz low-pass 4-pole Bessel filtering. Whole-oocyte ionic conductance was calculated by divided the current difference with voltage difference between –140 and –20 mV.

The concentration-relative conductance relationships for the GABA-induced inhibition (**Fig. 1e**) and ABA-induced activation (**Fig. 1g**) were fitted to a Hill equation using a least-squares method to derive the IC_50_ and EC_50_ values, the inhibitory concentration or the effective concentration, respectively, required for inducing a half maximal change^38,39^. The average of the normalized conductance were used to plot the data. All data are presented as the mean±s.e. (*n* = 12).

Methods for guard cell protoplast isolation and slow type anion current recording were described previously^36^. Before patch-clamp recording, 50 μM ABA ± 10 mM GABA were added to the bath solution for 40 min to modulate the S-type anion current.

### Stomatal aperture assays

Stomatal apertures were measured on epidermal peels from 3–4 week old plants grown in short-day conditions (120 μmol·m^-2^·s^-1^, 8 h light/16 h darkness, 22°C/18°C). Epidermal peels were incubated in opening buffer (0.1 mM CaCl, 50 mM KCl and 10 mM MES (2-Morpholinoethanesulphonic acid) with pH 6.15 by KOH) under light (200 μmol·m^-2^·s^-1^) for 2 h to induce stomatal opening. Epidermal peels were soaked in opening buffer with 10 μM ABA, 2 mM GABA, separately or in combination, or with an opening buffer under light control, for 2 h . Additionally for darkness assays in Fig S7, epidermal peels were soaked in opening buffer with or without 2 mM GABA in darkness or in light control for 2 h. The images were acquired using an inverted microscope (CX43; Olympus, Tokyo, Japan) at 40 × magnification, and stomatal apertures (width/length) measured with ImageJ software.

### GABA and ABA measurements

For GABA and ABA content measurements under drought and re-watering conditions, wild-type plants were sown on ½ Murashige and Skoog (MS) solid medium for 7 days and then transferred to pots containing 50 g (± 0.1 g) of 2:1 (w/w) soil:vermiculite and watered to weight. Two weeks, following the withdrawal of water, plants were collected since the soil water contents (SWC) dropped to 20% of the day “0” of the drought treatment. The well-watered plants were collected as control (CK). After 16 days, plants were re-watered and collected GABA and ABA extraction.

For GABA and ABA content measurements with ABA, saline, and osmotic treatments, After being sown on ½ MS solid medium for 7 days, seedlings were transferred onto ½ MS solid media containing 50 μM ABA, 120 mM NaCl, or 300 mM mannitol, and collected for GABA and ABA extraction About 100 mg of fresh plant material was ground under liquid nitrogen and incubated at 4°C for 10 h with ABA extraction buffer containing 80% methanol and 1% acetic acid (v/v) or GABA extraction buffer (0.2 mol L^-1^ hydrochloric acid). ABA and GABA content were detected by 1290 Ultra Performance Liquid Chromatography/6490 Triple Quadrupole Mass Spectrometer (Agilent, LC1290-6490) equipped with EclipsePlus C18 (2.1 mm × 100 mm, 1.8 μm). Solvents were ultrapure water with 0.01% acetic acid (A) and acetonitrile (B). ABA samples were eluted with a gradient from 20 to 80% solvent B, and GABA samples with 70% acetonitrile. Compounds are identified by retention time and mass/charge ratio.

### Drought assay

Wild-type, *gad2* mutant, and *GAD2*Δ *OE* lines were sown on ½ Murashige and Skoog (MS) solid medium for 7 days under short-day conditions (120 μmol·m^-2^·s^-1^; 8 h light/16 h dark) at 22°C. 7-day-old seedlings were transferred to pots containing 70 g (± 0.1 g) of 2:1 (w/w) soil:vermiculite, with 9 seedlings per pot. The pots were well-watered and randomly moved every day to minimise the impact of different environmental conditions. At 20 days after germination, seedlings were analysed as ‘Before drought’. Watering ceased for 4 weeks for *gad2* ‘Drought’ treatment and 5 weeks for *GAD2*Δ *OE*, with plants analysed at the end of this time. After rewatering for 2 days, plants were analysed as ‘Re-watered’.

### Transient expression in *Arabidopsis* protoplasts

Assays for transient expression in protoplasts were performed as described^40^. Expression vectors for *PYL1*, *ABI1*, *OST1*, and *ABF2* were obtained from Dr Yingfang Zhu (Henan University). *ProRD29A* was cloned into pGreenII 0800-LUC vector with primers listed in Supplementary Table 1. Transfected protoplasts were incubated for 12 h with 10 μM ABA and 1 mM GABA separately or in combination. Firefly luciferase (LUC) and Renilla luciferase (Ren) activities were detected by Multi-Mode Microplate Reader (Varioskan Flash).

### Real-time quantitative PCR (RT-qPCR) analysis

*Arabidopsis* seedlings were grown in ½ MS medium (with 1% (w/v) sucrose) for 7 days. Seedlings were transferred to ½ MS medium with or without 10 μM ABA and/or 2 mM GABA for 0.5, 1, or 3 h, at which point seedlings were harvested for total RNA extraction using the TRIzol reagent (Vazyme). The *gad* mutants *GAD2*Δ *OE* lines were treated with ABA for 1 h, and collected for RAN extraction. First-strand cDNA was synthesised from 1 mg total RNA with HiScript III RT SuperMix for qPCR (+gDNA wiper, Vazyme, R323-01). RT-qPCR was performed using ChamQ Universal SYBR qPCR Master Mix (Vazyme; Q711-02) on a 7500 Real Time PCR System (Applied Biosystems) following the manufacturer’s protocol. Transcript levels were calculated by normalisation to *Actin2/8* transcripts. Primers are listed in Supplementary Table 1.

### Seed germination and root elongation assays

For germination assays, *Arabidopsis* seeds were surface sterilised and stratified at 4°C for 72 h in the dark. About 100 seeds of every genotype were sown on the same plate containing ½ MS medium (with 1% (w/v) sucrose) with or without ABA, GABA, or muscimol at concentrations indicated in figures. The plates were kept at 22 °C under constant illumination (120 μmol·m^-2^·s^-1^; 16 h light/8 h darkness). Germination rate was defined as an obvious emergence of the radicle through the seed coat at the 5^th^ day, and the pictures were imaged at 7^th^ day. The germination rate assay and pictures imaging of *ABI1 OE* lines were completed at the 3^th^ and 6^th^ days respectively, for a rapid germination under ABA treatment.

For root elongation assays, *Arabidopsis* seeds were germinated as above for 2.5 days on ½ MS medium (with 1% (w/v) sucrose). Seedlings were then transferred to medium with or without ABA, GABA, or muscimol at the indicated concentrations and grown for 10 days. The primary root length was measured with ImageJ software.

### Pull-down assay

The Glutathione S-Transferase (GST) pull-down assay was performed as previously described^9^. Reaction mixtures containing 125 µg GST-ABI1 (amino acids 125–429), 25 µg His-PYL1 (amino acids 8–211) and resins (Glutathione Sepharose^TM^ 4B, Cytiva) with 10 μM ABA and 0, 0.01, 0.1, 1, and 10 mM GABA or 10 μM muscimol were incubated in pull down buffer containing 20 mM HEPES (2-[4-(2-hydroxyethyl) piperazin-1-yl] ethanesulfonic acid) pH 8.0, 150 mM NaCl, 1 mM TCEP (Tetrakis(2-cyanoethoxy)butane) and 5 mM MgCl_2_ for 12 h. Resins were washed ten times with washing buffer (50 mM Tris-HCl, pH 8.0), and remaining proteins collected by 50 mM reduced glutathione and separated by SDS-PAGE. The intensity of PYL1 bands were quantified by ImageJ software and standardised against the intensity of ABI1 in the ABA/no GABA control lane.

### Yeast two-hybrid assay

The coding sequence of *PYL1* and *ABI1* were cloned into pGADT7 and pGBKT7 vectors, respectively. Positive yeast transformants were selected on non-selective synthetic defined (SD)/-Leu-Trp medium. The interaction of PYL1 and ABI1 was tested at different ABA and GABA concentration on selective medium (SD/-Leu/-Trp/-His-Ade) at 30 °C for 4 d. Primers are listed in Supplementary Table 1.

### Split-luciferase and bimolecular fluorescence complementation (BiFC) assays

The split-luciferase assay was conducted as previously described^41^. Briefly, *ABI1-LUC^N^* and *PYL1-LUC^C^*with *35S:GFP* (*Green Fluorescent Protein*) or *35S:GAD2*Δ*-GFP* were co-expressed in *N. benthamiana* leaves for 2 days following *Agrobacterium*-mediated transient transformation. The leaves were sprayed with 50 μM ABA until the leaf surfaces were drenched to induce protein interaction. LUC signal intensity was captured by a cold charge-coupled device camera (CCD; Lumazone Pylon 2048B; Princeton), and fluorescence intensity analysed with ImageJ software.

BiFC assays were conducted as previously described^42^. Briefly, *ABI1-YFP^N^* and *PYL1-YFP^C^* were co-expressed with *35S:GAD2*Δ or pCAMBIA1300 vector control in *N. benthamiana* leaves for 2 days following *Agrobacterium*-mediated transient transformation. 50 μM ABA was sprayed onto the leaf surface to induce protein interaction. YFP fluorescence signals were captured by confocal laser scanning microscopy (Zeiss LSM 980 META) and the fluorescence intensity analysed with ImageJ software.

### MicroScale Thermophoresis (MST) analysis

MST assays were performed in a NanoTemper monolith NT.115. PYL1 and ABI1 proteins were concatenated with Cyan Fluorescent Protein (CFP) and His tags and purified as described^9^. Purified CFP-PYL1-His and CFP-ABI1-His proteins were concentrated and dialysed in PBS (Phosphate Buffer Solution) buffer (136 mM NaCl, 2.6 mM KCl, 10 mM Na_2_HPO_4_ and 2 mM KH_2_PO_4_ pH 7.4) with Amicon Ultra-15 centrifugal filter units (Millipore). Samples were mixed according to manufacturer’s instructions (NanoTemper). Binding assays were performed at 22 °C, 20% LED power, and 40% MST power. The *K_d_* value was calculated by Nano Temper Software MO affinity Analysis from triplicate experiments.

### Surface Plasmon Resonance (SPR) measurements

SPR assays were performed on Biacore T200 apparatus (GE Healthcare) with a CM5 sensor chip (Cytiva). Purified CFP-ABI1-His and CFP-His proteins were immobilised on CM5 sensor chip at 25°C. Serial dilutions of GABA (ranging from1.95 μM to 125 μM) in PBS-P (Phosphate-Buffered Saline with Tween®-20) were injected over the immobilized surface at a flow rate of 30 μL/min with a contact time of 120 s and a dissociation time of 300 s. All data was analysed by Biacore T200 Evaluation Software. The binding curve was fitted using the 1:1 Langmuir binding mode to calculate the *K_d_* value.

### *In vitro* phosphatase assay

OST1 autophosphorylation was used to analyse ABI1 phosphatase activity. For analyzing GABA affecting ABI1 activity, phosphorylation mixtures containing 6 µg His-ABI1, 2 µg MBP-OST1 with 10 mM GABA or 10 μM muscimol were incubated in kinase buffer (50 mM HEPES-MES pH 7.5, 1 mM MnCl_2_, 5 mM MgCl_2_, 1 mM DTT (Dithiothreitol), and 1 mM ATP-gamma-S (Adenosine 5′-[γ-thio]triphosphate tetralithium salt; Abcam, ab138911) for 30 min at 25 °C. For analyzing GABA affecting ABI1 activity with PYL1 and ABA-induction, phosphorylation mixtures containing 5 µg His-ABI1 (amino acids 125–429), 1 µg MBP-OST1, and 2 µg His-PYL1 with 10 µM ABA and 0, 0.01, 0.1, 1, and 10 mM GABA or 10 μM muscimol were incubated in kinase buffer for 30 min at 25 °C. *p*-Nitrobenzyl mesylate (Abcam, ab138910) was added to the reactions for 2 h at 25°C to tag phosphorylated proteins. Reactions were stopped by adding 10 µL SDS loading buffer and boiling for 5 min. The samples were separated by 10% SDS-PAGE. After electrophoresis, the proteins were transferred to PVDF membrane and immunoblotted with anti-thiophosphate ester antibody (Abcam, ab92570).

### *In vivo* phosphatase assay

*In vivo* phosphorylation status of OST1 was analysed as previously described^43^. Briefly, total proteins were extracted with IP buffer from 10-day-old *proSuper:OST1-GFP* plants treated with 2 mM GABA, 50 μM ABA, or both for 3 h. Proteins were separated on a 6% (w/v) phos-tag-PAGE gel containing 50 μM phos-tag acrylamide (AAL-107, Nard, Lot23H-02) and 50 mM MnCl_2_ (60 V, 5 h), or by 10% SDS–PAGE (120 V, 1.5 h). After electrophoresis, the phos-tag-PAGE gel was washed with buffer (10 mM EDTA (pH 8.0), 25.12 mM Tris, 191.82 mM glycine) 3 times for 10 min each, before proteins were transferred to nitrocellulose membranes at 100 V for 1 h. Phosphorylated and unphosphorylated OST1 proteins were detected by anti-GFP antibody (1:3,000). The Rubisco (rbcL) protein in SDS-PAGE served as the loading control. CIAP (Calf Intestine Alkaline Phosphatase) was involved as a negative control.

### Molecular docking and site-directed mutagenesis

Molecular docking between PYL1, ABI1 and GABA was performed using the Autodock Vina docking engine, executed with the AutodockTools-1.5.7 visualization software. The complex structure of PYL1–ABI1 was referred as previous studies^6,8,9^.

The method for site-directed mutagenesis of ABI1 has been described previously^36^. Briefly, the pGXBG-ABI1 construct was amplified using primers listed Supplementary Table 1. The amplified products were treated with *Dpn I* to digest the template, followed by transformation into *E. coli* DH5α. Clones containing the mutation were selected and used for two-electrode voltage clamp analysis to examine the antagonistic effect of GABA on ABA signalling.

### Quantification and statistical analysis

The number of biologically independent replicates for each experiment is indicated in figure legends. Statistical analyses were performed by one-way ANOVA, or ANOVA using Tukey’s comparison test.

## Acknowledgments

We thank *Prof.* Juan Xu (Zhejiang University) for providing *gad1*, *gad2*, and *gad1245* mutants; *Prof.* Zhi-Zhong Gong (China Agriculture University) for *abi1-3* mutant; *Prof.* Yang Zhao (CAS Center for Excellence in Molecular Plant Science) for *pyr1pyl12458* mutant; and *Prof.* Shu-Hua Yang (China Agriculture University) for *proSuper:OST1-GFP* line. We thank *Prof.* Ying-Fang Zhu (Henan University) for providing expression vectors for *PYL1*, *ABI1*, *OST1*, and *ABF2*; *Prof.* Matthew Gilliham and Natalie Betts (University of Adelaide) for project suggestion and writing support.

## Author contributions

Y.L. directed the projects. Y.L., C.-P.S., and conceptualization the research. Y.L., C.-Z.F., Y.-X.L., and H.Z. designed all the experiments. C.-Z.F., Y.-X.L., H.Z., P.L., and B.-R.L. performed the experiments and data analysis. Y.L., C.-P.S., Y.Z., and X.Z. discussed the project. Y.L. wrote the manuscript with the inputs from other authors. All authors have read, edited, and approved the content of the manuscript.

## Data, code, and materials availability

All data are available in the main text or the supplementary materials.

## Supplemental Figures

**Figure S1 Genotyping and drought assay of *gad* mutants and *GAD2***Δ ***OE* lines.**

**A.** Expression of *GAD2* in wild-type (WT) and transgenic *GAD2*Δ overexpression (OE) lines relative to expression of *Actin 2/8*. Data are mean ± s.e. (*n* = 3). **B.** GABA content in WT and *GAD2*Δ *OE* lines. Data are mean ± s.e. (*n* = 3). **C,D.** Representative images of WT and *gad2* (**C**) or *GAD2*Δ *OE* (**D**) plants before and after drought treatment and re-watering.

**Figure S2 GABA and ABA accumulation during saline and osmotic stress.**

**A,B.** ABA accumulation during osmotic (**A**) and saline (**B**) stress in wild-type *Arabidopsis*. **C, D.** GABA accumulation during osmotic (**C**) and saline (**D**) stress in wild-type *Arabidopsis*. **E.** ABA content in 7-day-old seedlings after GABA induction. **F**, Effect of *gad1245* mutation and *GAD2*Δ overexpression on ABA content under normal conditions and osmotic stress at 3 h after mannitol treatment. Data are mean ± s.e. (*n* = 3). FW, fresh weight.

**Figure S3 GABA blocks ABA inhibition of root elongation.**

**A.** Representative images of root growth of *Arabidopsis* seedlings exposed to different ABA, GABA, or muscimol combinations. **B.** Analysis of root growth shown in **a**. Box and whisker plots show interquartile range and extremes (*n* ≥ 60). Letters indicate significant differences at *P* < 0.05 by ANOVA with Tukey’s comparison test.

**Figure S4 GABA blocks the ABA-induced PYL1–ABI1 interaction.**

**A.** Yeast 2-hybrid assays testing PYL1 and ABI1 interaction at different ABA and GABA concentrations. **B.** Bifluorescence complementation assays using ABI1-YFP^N^ and PYL1-YFP^C^ in *N. benthamiana* leaves in wild-type (control) plants vs plants transformed with a *GAD2*Δ expression construct. GABA accumulation and fluorescence intensity are shown in ② and ③. Data are mean ± s.e. (*n* = 3). FW, fresh weight; a.u., arbitrary units.

**Figure S5 Binding specificity of GABA (muscimol) and ABI1.**

**A.** The binding curve between muscimol and CFP-ABI1, analysed by MST. Data are mean ± s.e. (*n* = 3). **B.** A representative SPR sensorgram of nitrate on the ABI1 chip from Figure 5E. **C.** The binding curve between GABA and CFP, analysed by SPR. **D.** A representative SPR sensorgram of nitrate on the CFP chip shown in **C**.

**Figure S6 GABA does not affect binding between PYL1 and ABA**, as analysed by MST.

**Figure S7 GABA blockage of dark-induced stomatal closure is not dependent on PYL1–ABI1.**

GABA inhibition of dark-induced stomatal closure in wild-type (WT) plants compared with *abi1* mutant (**A**), *ABI1 OE* (**B**), *pyr1pyl12458* sextuple mutant (**C**), *PYL1 OE* (**D**), and *almt12* mutant (**E**) lines. Box and whisker plots show interquartile range and extremes (*n* > 120). Letters indicate significant differences at *P* < 0.05 by ANOVA with Tukey’s comparison test. Percentages indicate the GABA caused stomatal closure decreasing compared with dark treated control.

**Figure S8. GABA blockage of ABA-induced germination inhibition is ABI1- and PYL1-dependent.**

**A.** Expression of *ABI1* in wild-type (WT) and transgenic *ABI1* overexpression (OE) lines relative to expression of *Actin 2/8*. Data are mean ± s.e. (*n* = 3). **B.** Expression of *PYL1* in WT and *PYL1* OE lines, relative to expression of *Actin 2/8*. Data are mean ± s.e. (*n* = 3). **C–J**, Seed germination response to ABA inhibition in *abi1* mutant (**C** and **D**), *ABI1 OE* lines (**E and F**), *pyr1pyl12458* sextuple mutant **(G and H**), and *PYL1 OE* lines (**I and J**). For **D**, **F, H**, and **J**, data are mean ± s.e. (*n* > 100). Letters indicate significant differences at *P* < 0.05 by by one-way ANOVA test.

**Figure S9 GABA does not bind to the ordered region of ABI1, as analysed by MST.**

**Table S1 Primer list**

