## supplemental figures for "GABA antagonises ABA perception and signalling in plants"

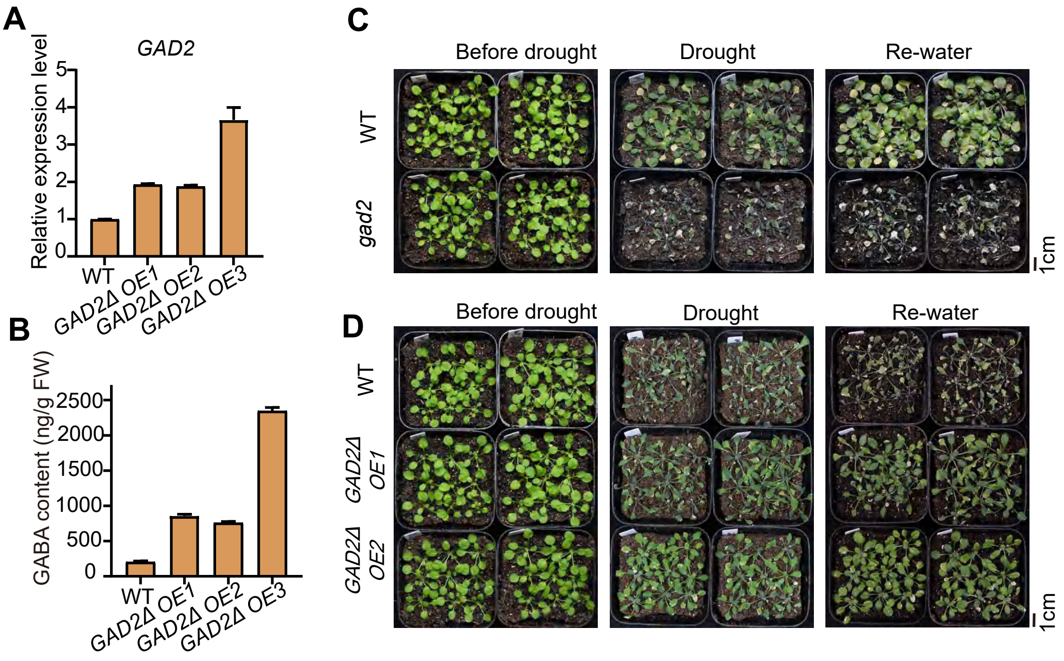


**Figure S1 Genotyping and drought assay of *gad* mutants and *GAD2Δ OE* lines.**
**A.** Expression of *GAD2* in wild-type (WT) and transgenic *GAD2Δ* overexpression (OE) lines relative to expression of *Actin 2/8*. Data are mean ± s.e. ($n = 3$). **B.** GABA content in WT and *GAD2Δ OE* lines. Data are mean ± s.e. ($n = 3$). **C,D.** Representative images of WT and *gad2* (**C**) or *GAD2Δ OE* (**D**) plants before and after drought treatment and re-watering.

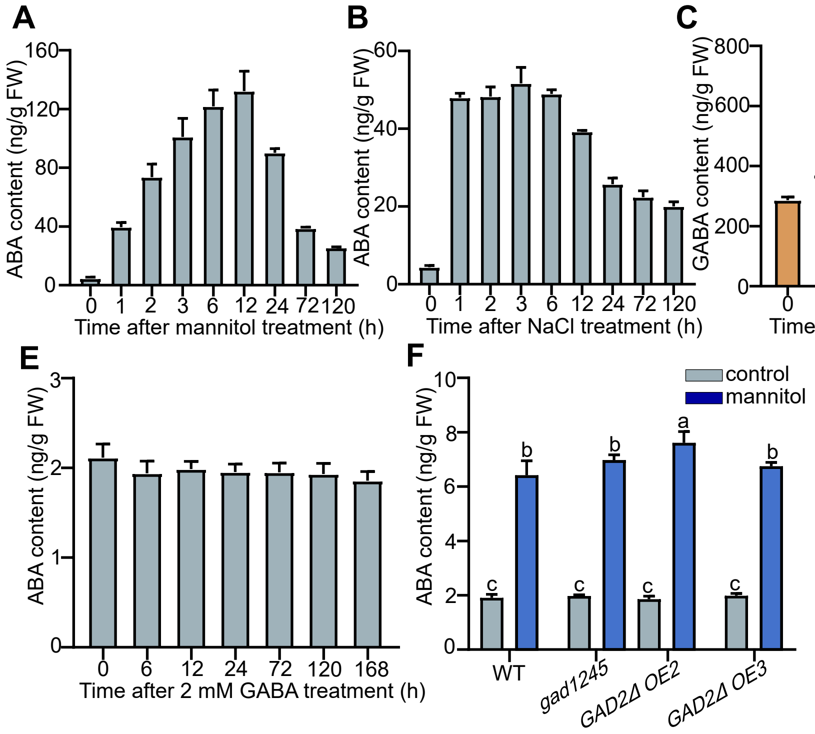


**Figure S2 GABA and ABA accumulation during saline and osmotic stress.**
**A,B.** ABA accumulation during osmotic (**A**) and saline (**B**) stress in wild-type *Arabidopsis*. **C, D.** GABA accumulation during osmotic (**C**) and saline (**D**) stress in wild-type *Arabidopsis*. **E.** ABA content in 7-day-old seedlings after GABA induction. **F**, Effect of *gad1245* mutation and *GAD2Δ* overexpression on ABA content under normal conditions and osmotic stress at 3 h after mannitol treatment. Data are mean ± s.e. ($n$ = 3). FW, fresh weight.

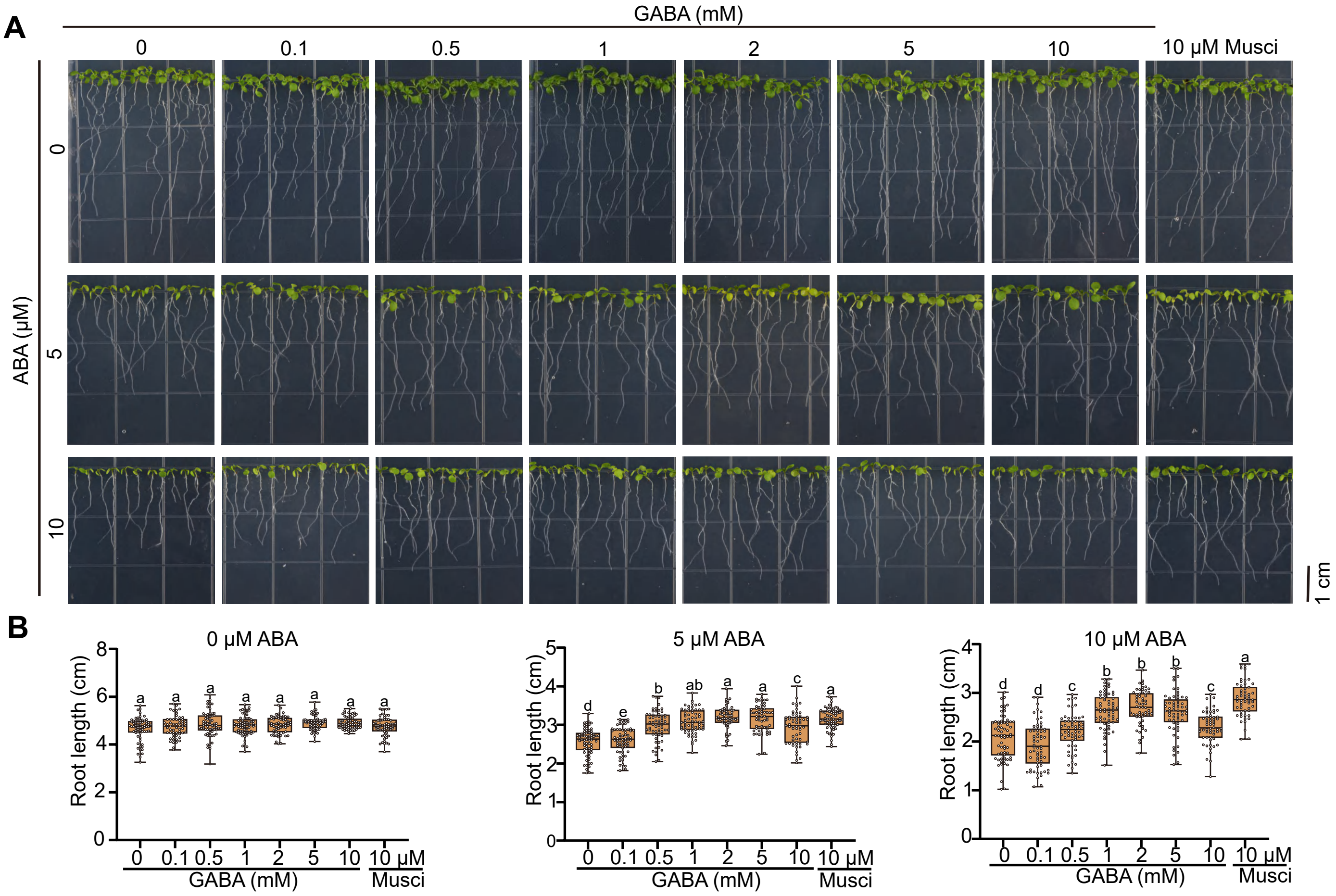


**Figure S3 GABA blocks ABA inhibition of root elongation.**

**A.** Representative images of root growth of *Arabidopsis* seedlings exposed to different ABA, GABA, or muscimol combinations. **B.** Analysis of root growth shown in **a**. Box and whisker plots show interquartile range and extremes ($n \geq 60$). Letters indicate significant differences at $P < 0.05$ by ANOVA with Tukey ’ s comparison test.

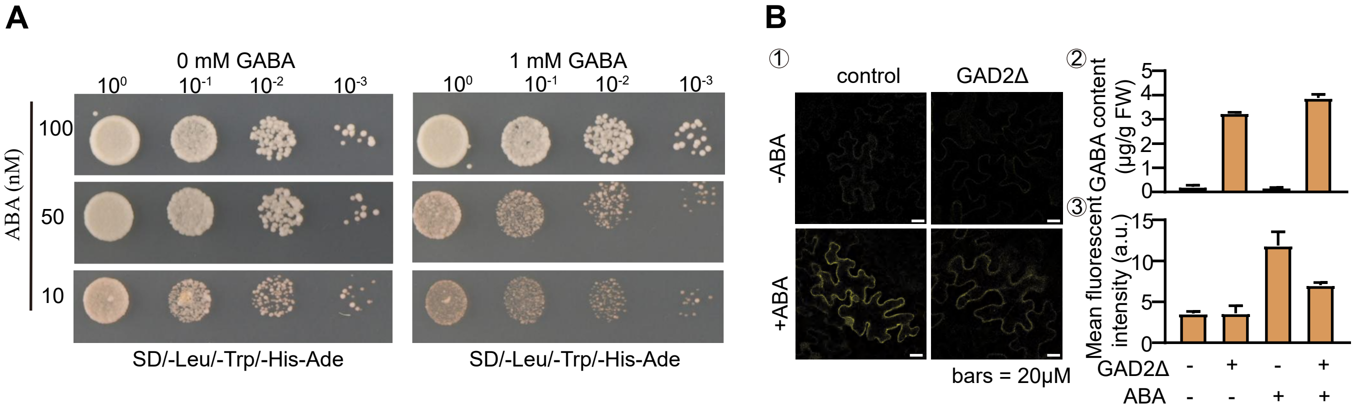


**Figure S4 GABA blocks the ABA-induced PYL1 – ABI1 interaction.**

**A.** Yeast 2-hybrid assays testing PYL1 and ABI1 interaction at different ABA and GABA concentrations . **B.** Bifluorescence complementation assays using ABI1-YFP$^{N}$ and PYL1-YFP$^{C}$ in *N. benthamiana* leaves in wild-type (control) plants vs plants transformed with a *GAD2Δ* expression construct. GABA accumulation and fluorescence intensity are shown in k and l . Data are mean ± s.e. (*n* = 3). FW, fresh weight; a.u., arbitrary units.

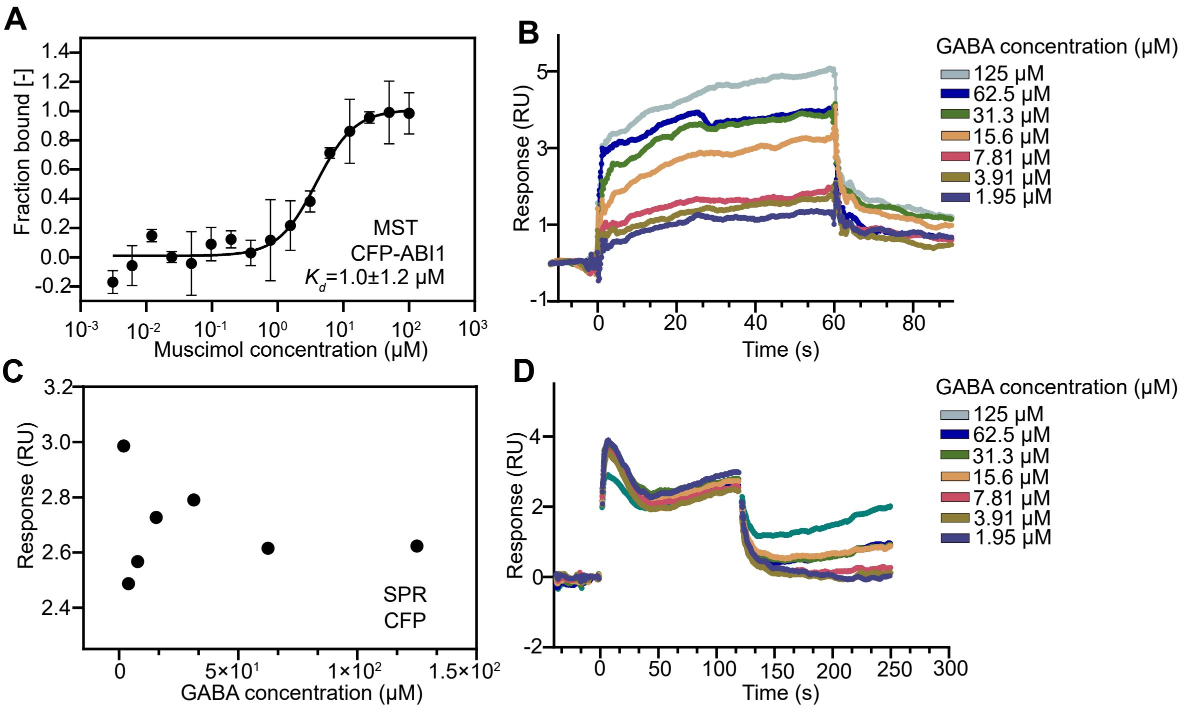


**Figure S5 Binding specificity of GABA (muscimol) and ABI1.**
**A.** The binding curve between muscimol and CFP-ABI1, analysed by MST. Data are mean ± s.e. ($n$ = 3). **B.** A representative SPR sensorgram of nitrate on the ABI1 chip from **Figure 5E**. **C.** The binding curve between GABA and CFP, analysed by SPR. **D.** A representative SPR sensorgram of nitrate on the CFP chip shown in **C**.

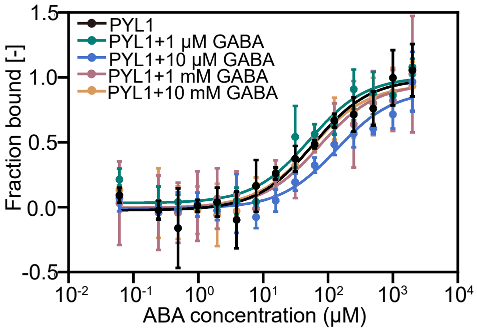


**Figure S6 GABA does not affect binding between PYL1 and ABA**, as analysed by MST.

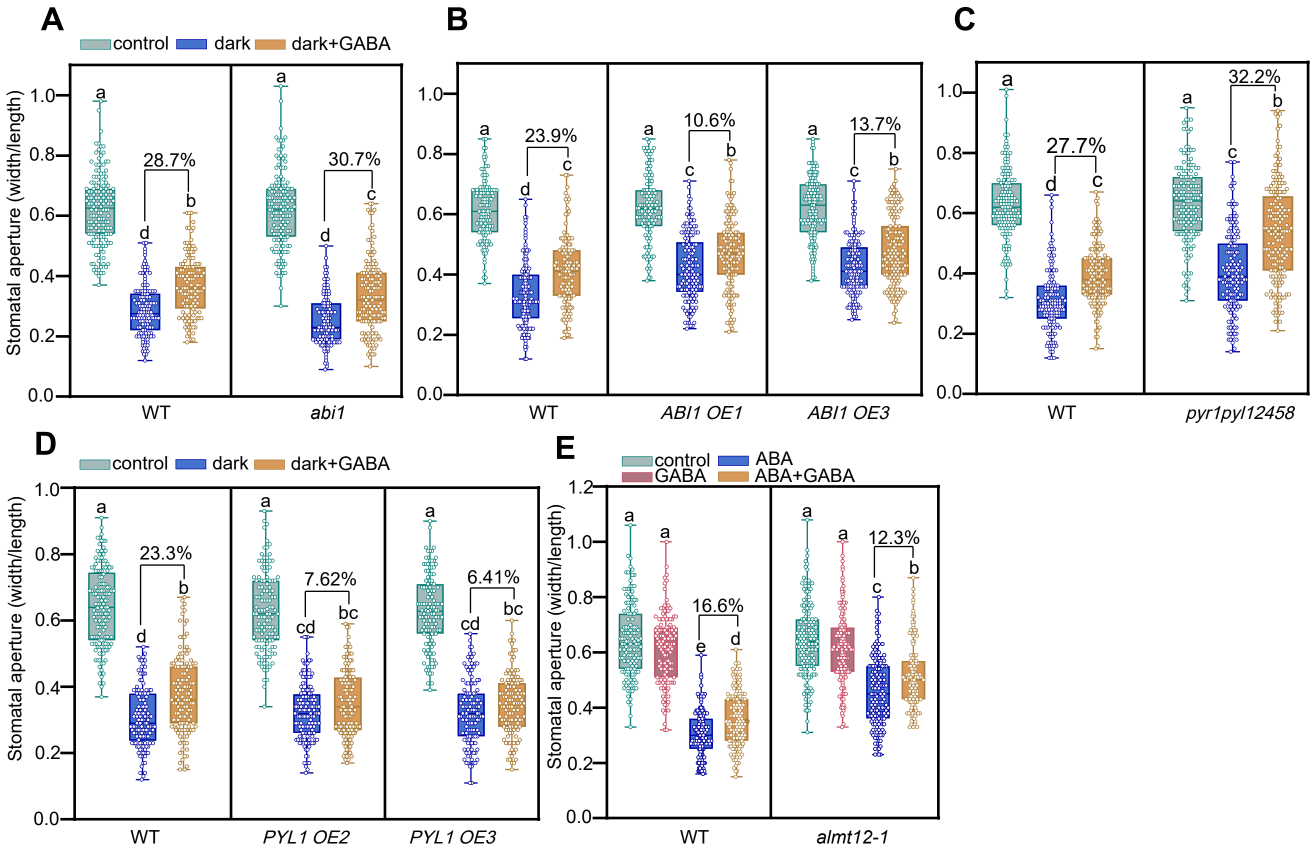


**Figure S7 GABA blockage of dark-induced stomatal closure is not dependent on PYL1 – ABI1.**
GABA inhibition of dark-induced stomatal closure in wild-type (WT) plants compared with *abi1* mutant (**A**), *ABI1 OE* (**B**), *pyr1pyl12458* sextuple mutant (**C**), *PYL1 OE* (**D**), and *almt12* mutant (**E**) lines. Box and whisker plots show interquartile range and extremes ($n$ > 120). Letters indicate significant differences at $P$ < 0. 05 by ANOVA with Tukey ’ s comparison test. Percentages indicate the GABA caused stomatal closure decreasing compared with dark treated control.

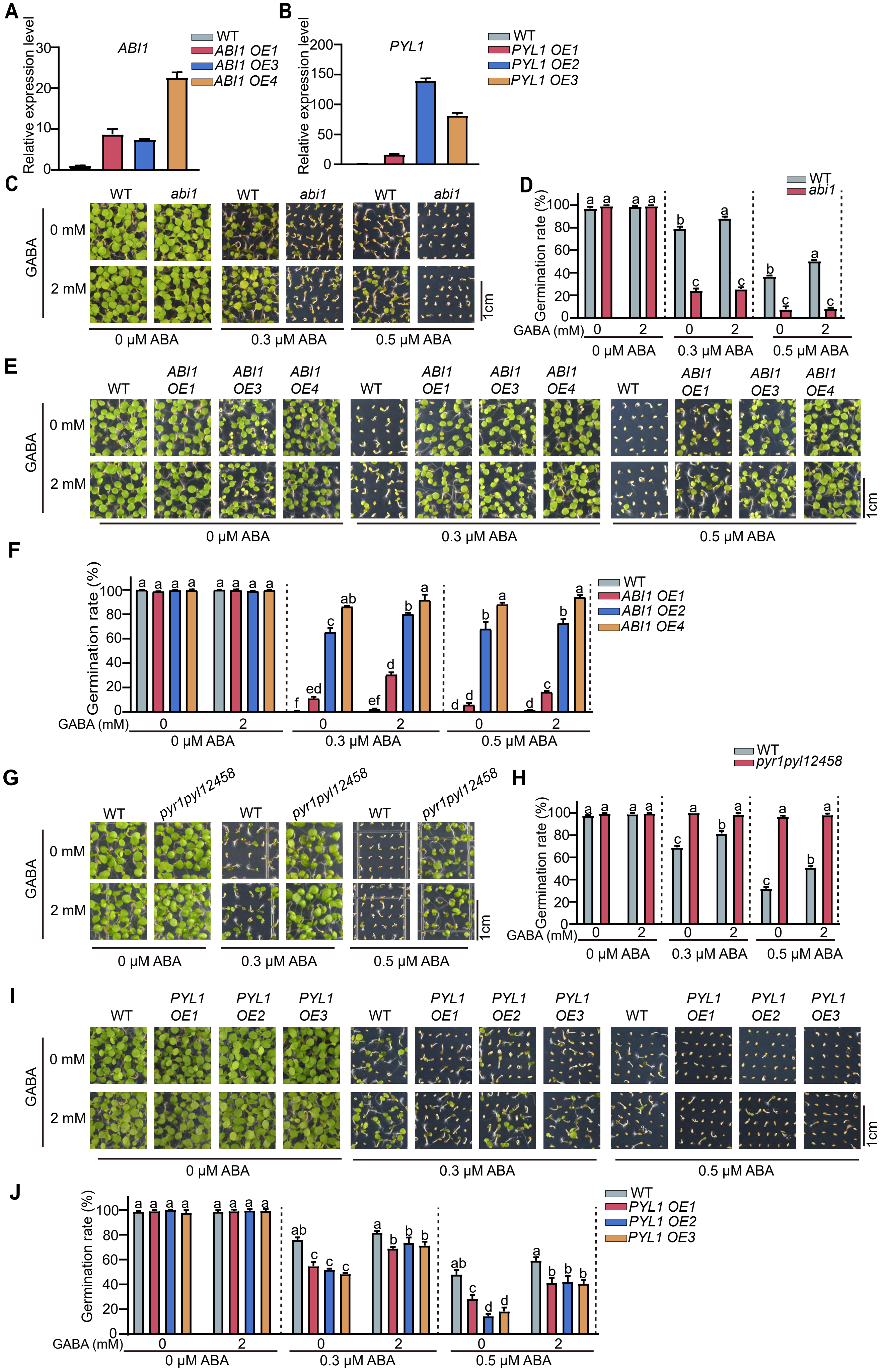

A
ABI1
Relative expression level
WT
ABI1 OE1
ABI1 OE3
ABI1 OE4
B
PYL1
WT
PYL1 OE1
PYL1 OE2
PYL1 OE3
C
D
WT
abi1
Germination rate (%)
GABA (mM)
0 μM ABA
0.3 μM ABA
0.5 μM ABA
GABA
0 mM
2 mM
1cm
E
F
ABI1 OE2
G
H
pyr1pyl12458
I
J

**Figure S8 GABA blockage of ABA-induced germination inhibition is ABI1- and PYL1-dependent.**
**A.** Expression of *ABI1* in wild-type (WT) and transgenic *ABI1* overexpression (OE) lines relative to expression of *Actin 2/8*. Data are mean ± s.e. ($n$ = 3). **B.** Expression of *PYL1* in WT and *PYL1* OE lines, relative to expression of *Actin 2/8*. Data are mean ± s.e. ($n$ = 3). **C – J**, Seed germination response to ABA inhibition in *abi1* mutant (**C** and **D**), *ABI1 OE* lines (**E and F**), *pyr1pyl12458* sextuple mutant **(G and H**), and *PYL1 OE* lines (**I and J**). For **D**, **F, H**, and **J**, data are mean ± s.e. ($n$ > 100). Letters indicate significant differences at $P < 0.05$ by by one-way ANOVA test.

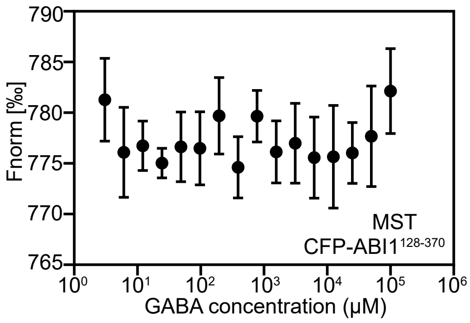


**Figure S9 GABA does not bind to the ordered region of ABI1,** as analysed by MST.
